# FEABAS: A Stitching and Alignment Tool for Serial EM Data

**DOI:** 10.64898/2026.06.07.730510

**Authors:** Yuelong Wu, Jeff W. Lichtman

**Affiliations:** Department of Molecular and Cellular Biology, Center for Brain Science, Harvard University, Cambridge, USA

## Abstract

Volume electron microscopy (vEM) is the most advanced and scalable technique for reconstructing synaptic-level wiring diagrams of the nervous system. Following image acquisition, the first critical step is reconstruction of a digitized volume, which assembles millions of electron microscope images into a coherent 3D volume that will underpin all downstream analyses. Existing methods work best with artifact-free datasets or rely on computationally intensive deep learning approaches or time-consuming human editing, restricting the broader applicability of vEM. To circumvent these challenges, we have developed FEABAS, a scalable, cross-platform, open-source software package designed to elastically montage and align electron microscope image datasets with high efficiency and precision. It leverages adaptive mesh modeling and finite element methods, enabling robust handling of datasets containing common artifacts such as wrinkles, folds, tears, and broken sections, while maintaining a lightweight, accessible implementation suitable for diverse computational environments.

## Introduction

Volume electron microscopy (vEM) remains the most established technology for reconstructing neuronal circuits at the synapse level, offering unparalleled detail and clarity of microstructures at nanometer resolution. The core principle of vEM involves sequential imaging of a heavy metal-stained specimen, slice-by-slice at nanometer z-step increments, with slicing achieved either by serial sectioning of the specimen (ssEM) or block-face imaging (e.g. SBF-SEM, FIB-SEM, GCIB-SEM). The limited field of view of the high-resolution electron microscopes further requires the acquisition of numerous overlapping image tiles per slice. As a result, a minuscule volume of 1mm^3^ could easily yield more than 100 million image tiles. Conceptually, volume reconstruction reverses this imaging process in silico: 2D stitching mosaics the tiles into slices, and subsequent 3D alignment transforms the slices into a continuous, coherent volumetric dataset. While this reconstruction serves merely as a technical vehicle for biological discovery, its accuracy is foundational. Even subtle local misregistration can critically undermine the subsequent step of automatic segmentation, thereby compromising the fidelity of downstream analysis.

Numerous computational solutions exist for vEM volume reconstruction. Most software packages are focused on either 2D stitching or 3D alignment, with few solutions offering an integrated workflow. For 2D stitching, methods generally rely on sparse feature matching to identify correspondences between adjacent tiles, and subsequently minimize registration errors by calculating transformation models ranging from rigid to non-linear spline-based functions. Similarly, 3D alignment often extends this matching-to-transformation logic to the z-axis, often modeling the volume as a connected spring-mesh system that undergoes global relaxation to minimize registration errors. These classical approaches^1–6^ frequently suffer from mechanical oversimplification, lacking the flexibility to robustly correct the high-order deformations and complex artifacts inherent to challenging vEM datasets. While deep learning (DL) approaches have recently emerged^7–9^ as competitive alternatives, they introduce significant training and computational overheads, often demanding specialized expertise to implement effectively.

Here we present FEABAS (Finite-Element Assisted Brain Assembly System), a fully integrated, open-source Python workflow for both elastic 2D stitching and 3D alignment, designed for easy deployment across diverse platforms and scalable environments. While rooted in the principles of classical spring-mesh systems, FEABAS employs a Finite Element analysis (FEA) formulation to model tissue deformation with greater flexibility and efficiency. By utilizing adaptive mesh and robust sparse linear solvers, the software efficiently handles complex non-linear distortions, including severe wrinkle artifacts. Crucially, for the 3D alignment, the workflow prioritizes a robust thumbnail-resolution alignment step, ensuring global coherence before high-resolution optimization. FEABAS is designed for immediate, out-of-the-box utility on well-conditioned datasets; and for challenging volumes, it offers advanced options that yield reconstruction quality comparable to deep learning pipelines, without the associated training overhead or hardware constraints. FEABAS is implemented entirely for CPU execution, ensuring its utility across diverse computing environments without the need for GPU-specific hardware or drivers.

## Methods

### Integrated vEM Reconstruction Workflow

FEABAS implements a fully integrated, three-stage workflow: 2D Stitching, Thumbnail Alignment, and Fine Alignment (Figure 1). Starting from raw tiles, the 2D Stitching stage mosaics and renders full-resolution stitched sections. These are immediately downsampled for Thumbnail Alignment, which computes a coarse global registration. This coarse solution initializes the Fine Alignment stage, where grid-based template matching is performed on the full-resolution sections (pre-transformed by the coarse results) to resolve intricate local deformations. All three stages employ a unified algorithmic foundation: corresponding landmarks found by matching image patches sparsely drives a Finite Element Analysis (FEA) relaxation on adaptive meshes, minimizing elastic energy to generate the transformation fields that register the images.

**Figure 1:**
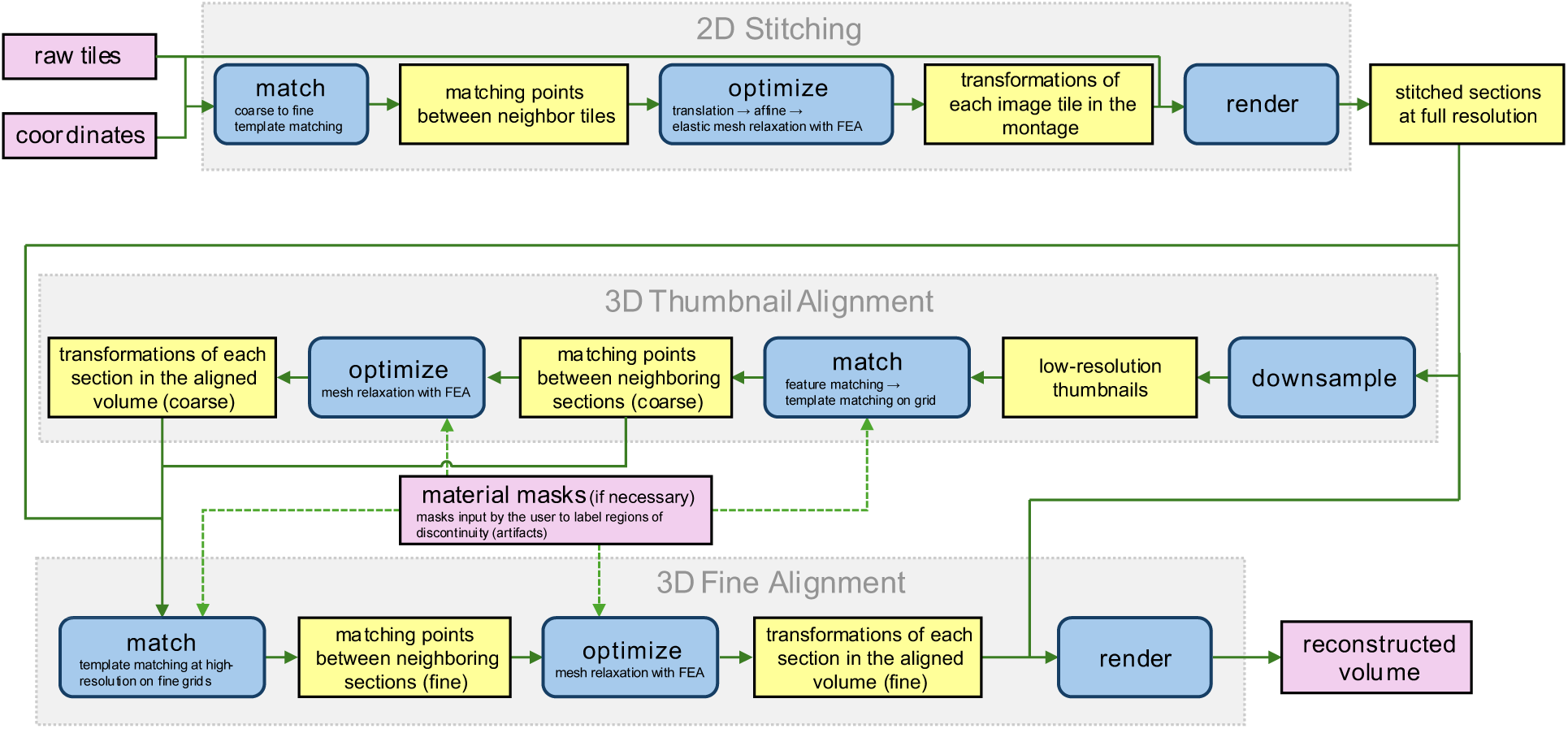
The FEABAS Workflow for Automated vEM Reconstruction. Reconstruction is divided into three interleaved phases: (1) 2D Stitching of individual image tiles, (2) Thumbnail Alignment for global Z-axis continuity, and (3) Fine Volumetric Alignment for localized feature registration. The output from upstream phases gives the input or initialization for downstream operations (arrows).

### Algorithmic Foundation

#### Mesh Generation

FEABAS represents image transformations as displacement fields piecewise linearly mapped onto a discretized domain; as such, mesh generation serves as the foundation for the entire workflow. To achieve an adaptive discretization, FEABAS utilizes Planar Straight-Line Graphs (PSLGs) as the geometric basis for mesh construction. These graphs define piecewise-linear boundaries that partition the domain into closed polygons, each representing a region with specified mesh constraints, including target element size and minimum interior angles, and assigned mechanical properties such as stiffness. Following PSLG initialization, a Delaunay refinement algorithm^10^ generates high-quality triangular meshes that strictly respect these internal boundaries.

This framework allows for targeted computational efficiency and modeling flexibility. For example, during 2D stitching, FEABAS automatically generates higher mesh densities along tile overlaps to concentrate degrees of freedom where seam errors may rise and need to be corrected. For 3D alignment, the software defaults to a uniform mesh over artifact-free image areas but supports advanced customization via material masks. These color-coded masks, which can be generated manually or through deep learning segmentation^11^, allow users to assign heterogeneous mesh geometries and elastic constants to specific biological regions or artifacts, ensuring that the model’s physical response is tuned to the sample’s underlying heterogeneity.

#### Image Matching

FEABAS employs two types of matching strategies to generate the displacement constraints required for mesh relaxation. The choice between feature-based and template-based matching depends on the expected degree of geometric distortion: feature matching is prioritized for coarse alignment and large rotations, while template matching is used for high-precision local refinement.

For stitching or local registration, we perform cross-correlation in the Fourier domain to maximize computational throughput. To enhance robustness, a band-pass filter is applied to the spectral representations, which suppresses both high-frequency noise and low-frequency intensity gradients, thereby isolating the structural textures most relevant to registration. To reject spurious correspondences due to ambiguous correlation profiles, we implement a peak-to-null validation procedure. Each template is matched against its spatially flipped matching partner to establish a baseline noise floor. A match is only considered high confidence if the primary correlation peak exceeds this “flipped” peak by an empirical threshold.

For scenarios involving large-scale rotations or significant non-linear deformations, such as thumbnail alignment, FEABAS utilizes a specialized feature-matching pipeline. Keypoints are localized as extrema in images processed with a Difference-of-Gaussian (DoG) filter. Unlike standard methods such as SIFT, our implementation omits the scale-space pyramid, optimizing the detector for the fixed-resolution structural landmarks inherent to vEM data. To ensure rotation invariance, we generate descriptors based on the Radon transform of local image patches centered on each keypoint. The resulting sinogram descriptors represent the structural projection of the patch across multiple angles. The rationale for this design is that the Radon transform, as an integral operation, provides superior robustness to the stochastic noise and the diverse range of artifacts typical of EM images compared to gradient-based descriptors. Similarity between two keypoints is computed as the normalized cross-correlation (NCC) of their sinogram descriptors along the angular dimension. By identifying the maximum correlation value across all possible angular shifts, the descriptor remains inherently robust to arbitrary image rotations. Feature point correspondences are established through an exhaustive similarity search, where each keypoint is assigned with its most similar counterpart in the target image. Matches are subsequently refined for spatial consistency based on the geometric criteria outlined in the Hierarchical Iterative Matching and Relaxation section.

#### Finite Element Formulation and Elastic Relaxation

FEABAS computes transformations by minimizing the total potential energy of the discretized spring-mesh system, effectively bridging discrete image matches (springs) and continuous geometric deformations (meshes) while enforcing physical plausibility constraints. We represent the state of the mesh using a global displacement vector ***u*** ∈ ℝ^2n^, which concatenates the x and y displacements of all n nodes. We employ constant-strain triangle (CST) elements under a linear elasticity model, where the elastic energy of the mesh is a quadratic function of ***u***.

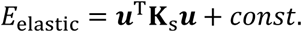

The global stiffness matrix **K**_s_ is constructed by assembling individual element-wise matrices ***k***_*e*_. For each element, ***k***_*e*_ is derived based on its local geometry (area and aspect ratio) and its material properties (Young’s modulus and Poisson’s ratio). This shape-dependent formulation ensures that the physical response is not sensitive to the discretization pattern. These local matrices are mapped to their corresponding global indices and summed to form the sparse global matrix **K**_s_.

To incorporate registration data, we model each matched point pair as a zero-resting-length spring. For a given match at a specific location, the displacement is interpolated from the nodes of its enclosing mesh element using barycentric coordinates. This mapping allows expression of the matching potential energy in the same quadratic form as the elastic energy:

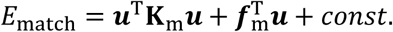

where **K**_m_ represents the spring stiffness assigned to the matches and ***f***_m_ is the load vector derived from the initial displacement error between matched points.

The optimal transformation is found by minimizing the total potential energy of the system, *E*_total_, defined as the weighted sum of the elastic strain energy and the matching constraint energy:

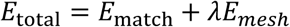

where *λ* is a scalar weighting factor that balances the relative contribution of the elastic resistance from the meshes against the driving forces from the matching springs. To ensure stable and balanced relaxation, FEABAS defaults to a trace-normalization strategy, where *λ* is automatically determined such that:

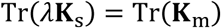

By equalizing the traces of the stiffness matrices, the framework ensures that the “global stiffness” of the matching constraints is commensurate with the mesh’s internal elastic resistance, regardless of the number of matches or the mesh density.

Taking the derivative of the total energy with respect to the displacement vector ***u*** and setting it to zero yields the governing equilibrium equation:

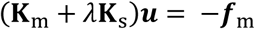

This sparse linear system is then solved to find the displacement field that minimizes the global error while preserving the model’s physical plausibility.

To accelerate convergence and ensure the solver begins within the optimal basin of attraction, FEABAS performs a global rigid initialization before assembling the full FEA system. For each mesh, an initial transformation is estimated by fitting the available matching points in a Procrustes-style alignment step. After the global rigid initialization and obtaining the governing equation, FEABAS uses the Minimal Residual (MINRES) algorithm as its default solver. To accelerate convergence, two types of preconditioning strategies are used. For standard registration tasks, a Jacobi preconditioner is used. In scenarios involving large deformations, a smoothed aggregation algebraic multigrid (AMG)^12^ preconditioner is employed for its effectiveness in dealing with low-frequency errors. Solver termination is governed by two complementary convergence criteria. First, the process stops if the relative residual falls below a predefined tolerance. Second, we implement an energy-based exit strategy: the solver terminates if the energy reduction between iterations is smaller than the expected energy fluctuation caused by a sub-pixel perturbation (e.g., 0.1 pixels) of the nodal locations. This criterion ensures that computational effort is only expended when it yields physically significant improvements to the alignment.

For standard registration, which yields a large but sparse linear equation, this system is solved directly. To handle physical non-linearities, such as modeling wrinkles that are “easy to expand but difficult to compress”, FEABAS utilizes the iterative Newton’s method. In each iteration, the element-wise constitutive matrices are updated based on the current strain state, and the global system is re-assembled and re-solved until convergence or reaching a predefined maximum iteration number.

In some applications, for example, when registering new images to an existing aligned volume, FEABAS allows designating specific meshes as reference sections (anchors). These anchors are enforced by imposing Dirichlet boundary conditions on the nodal displacements. Mathematically, this is achieved through an efficient system reduction: the rows and columns corresponding to the fixed degrees of freedom (DOFs) are removed from the global stiffness matrix and the load vector prior to solving.

Note that the linear elastic stiffness matrix employed in our standard formulation is based on the infinitesimal strain assumption, which is not frame-invariant under large finite rotations. While this assumption holds for most stitching applications where angular offsets are minimal, it can introduce artifacts during 3D alignment if sections are significantly misoriented before FEA formulation. To mitigate this, we perform a global rigid pre-alignment of all meshes to a common reference frame prior to constructing the FEA system. In rare instances of severe intra-section distortion where local rotations exceed the linear regime, we have implemented an optional St. Venant–Kirchhoff hyperelastic model^13^. This slightly more complex formulation provides a rotation-invariant energy functional at the cost of moving to a non-linear optimization framework. However, in our experience across the datasets presented here, the linear model combined with rigid pre-transformation proved sufficient for high-fidelity reconstruction, and the hyperelastic implementation remains an infrequently utilized feature for theoretical completeness and potential extreme cases (Supplementary Figure 1).

#### Hierarchical Iterative Matching and Relaxation

To achieve efficient registration across varying scales, FEABAS employs a hierarchical feedback loop between image matching and FEA relaxation. The process begins with a coarse matching phase, typically using feature-based descriptors or large-window template matching, to estimate an initial displacement field, which then serves as a spatial prior for subsequent iterations. By dynamically re-centering and constraining search windows based on the current mesh deformation, the system progressively refines the registration. By iteratively updating the matching constraints, the system can identify and prune outlier matches that contradict the physical elasticity of the surrounding mesh, making the process more robust in regions of low image quality or significant artifacts.

#### Image Rendering

Throughout the FEABAS pipeline, the rendering engine serves as a critical utility for both intermediate registration cycles and final volume synthesis. To map pixels from the source image to the target coordinate space represented by the mesh displacement, we first employ a trapezoidal map algorithm^14^ for efficient point-location, identifying the specific triangular mesh element associated with each output pixel. Within each element, the local displacement field is calculated via piecewise linear interpolation of the nodal displacement vectors. Image resampling is performed using Lanczos interpolation ^15^, a high-order filter that preserves structural detail and minimizes aliasing artifacts during the transformation.

In practice, inaccuracies in material masks or stochastic errors in the matching process can occasionally result in localized mesh self-intersections. Due to the technical limitation of the standard implementations^16^ of trapezoidal map algorithm that require a valid, non-self-intersecting planar graph, FEABAS includes a dedicated resolution layer to ensure algorithmic stability. When conflicting regions are identified and localized, the rendering engine partitions them along the mid-line of the collided area and renders each part based on their proximity to the colliding boundaries. The software can also utilize semantic rendering priorities defined within the grayscale material masks to override the mid-line partitioning strategy. This allows the system to intelligently manage overlaps: regions labeled as high-priority (e.g., intact neural tissue) are preserved over low-priority regions (e.g., resin-filled cracks or substrate artifacts).

This ensures that throughout the rendering process, the most biologically relevant data remains visible.

### Pipeline Implementation

#### 2D Stitching

The first stage of the reconstruction process is 2D stitching, where individual image tiles are assembled into a seamless 2D mosaic. The stitching pipeline in FEABAS does not rely on predefined grid assumptions. Instead, it utilizes the raw spatial metadata, including image paths, tile dimensions, and estimated stage coordinates, to automatically reconstruct the neighborhood topology and identify all overlapping tile pairs. This flexibility allows FEABAS to handle diverse scanning geometries, ranging from standard rectilinear grids to more complex patterns like the hexagonal arrangements used by Zeiss MultiSEM systems^17^.

Once overlapping tile pairs are identified, FEABAS establishes correspondences through a multi-stage matching strategy designed to overcome image artifacts and initial coordinate inaccuracies. For each overlapping pair, the pipeline crops out the potential overlap regions and performs a global template matching pass to estimate the general translation. To handle cases where severe artifacts might obscure the matching signal, the system employs a fallback strategy: if the global match confidence falls below a threshold, the overlap is subdivided into smaller blocks for localized matching; the sub-block pair yielding the highest confidence score is then selected to represent the transformation. This local search increases the likelihood of identifying high-confidence features in clean sub-regions. Following the global translation estimate, the rectangular overlapping areas are meshed for both tiles of a pair. The system then initiates the hierarchical matching and relaxation iterations described in the algorithmic foundation section, beginning with a coarse grid and progressively increasing grid density, alternating between template matching and FEM relaxation. Upon completion, the final correspondences are saved alongside the mesh elastic energy required to achieve the match. Saving this energy metric allows the downstream transformation optimizer to identify tiles with significant local distortions (e.g., from scanning charging effects) and automatically assign them a lower elastic modulus (softer mesh) in the final relaxation to prevent localized errors from propagating through the mosaic.

Recognizing the occasional inaccuracies in the initial stage metadata, FEABAS re-estimates the global positions of all tiles simultaneously using a linear least squares (LLS) optimization with the successful matches as constraints. This corrected arrangement is then used to re-calculate search windows for previously failed matches and attempt matching a second time, allowing the system to recapture matches that were previously missed due to metadata inaccuracies.

The optimization phase integrates individual tile correspondences into a unified 2D mosaic. The optimization solver begins by estimating the relative translations of all tiles through Linear Least Squares (LLS). Following initialization, the system constructs an FEA mesh for each tile. Employing a spatially adaptive density strategy, meshes are refined at overlapping boundaries to capture high-frequency deformations while remaining coarse in non-overlapping regions to maximize computational efficiency.

To account for systemic distortions, such as lens distortion or ones caused by the shared optical path in MultiSEM^17^, FEABAS offers an optional intermediate relaxation step, group elastic relaxation, where groups of tiles are constrained to share a common deformation field. This is achieved through a coordinate transformation that reduces the system’s degrees of freedom (DOFs). By introducing a reduced displacement vector ***q*** that contains only the unique DOFs after displacement sharing, and a sparse binary matrix **S** that maps the reduced set of unique DOFs to the full set of nodal displacements, we express the relationship as ***u*** = **S*q*.** Substituting this into the energy functional and taking the derivative yields the reduced system equation:

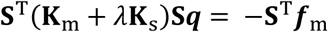

Solving for *q* provides the optimal collective transformation for the constrained group. These values are then mapped back to the original nodes, and the resulting coordinates are established as the new resting state of the meshes for the following relaxation steps. This effectively bakes in the systemic corrections before the final refinement.

Once the shared deformations are resolved, the constraints are lifted to allow for a final independent stitching relaxation. In this step, each tile mesh is permitted to deform freely to resolve any remaining localized misregistration. To ensure robustness against false correspondences, the solver employs iterative residual pruning: matching points exhibiting large residual error at an intermediate convergence stage are identified as erroneous and filtered out.

In instances where the specimen contains disconnected regions, such as discrete tissue patches separated by featureless resin, the lack of correspondences can result in a fragmented tile graph. To maintain a globally consistent mosaic, FEABAS performs a connected component analysis on the finalized matching graph. If multiple isolated clusters are identified, the system uses the initial-stage coordinates at the interfaces between these components to calculate their relative spatial offsets. Each component is then subjected to a rigid translation to its metadata-derived position. This anchoring strategy effectively bridges gaps where image features are absent, ensuring the global section geometry is preserved without biasing or introducing artificial matches into the regions supported by high-confidence physical matches.

The final step of the stitching pipeline involves rendering the individual tiles into a unified 2D mosaic using the image rendering routine described previously. To ensure a seamless visual result that is optimized for subsequent 3D alignment, FEABAS employs a global intensity equalization step. During the matching step, the mean and standard deviation of pixel intensities are calculated for all overlapping regions. These paired statistics are used to formulate a regularized least-squares problem, solving for tile-specific brightness offsets using the means, contrast multipliers using the logarithm of the standard deviation. Because an overlapping region is scanned first as a virgin surface on one tile and subsequently as a pre-exposed surface on the neighboring tile, there are subtle differences in image intensities. An *L_2_* regularization term is included to prevent such localized intensity variations from accumulating into systematic global intensity gradient. Upon convergence, the optimized parameters are linearly applied to each tiles. The blending at the tile interfaces follows a frequency-based blending strategy. Images are decomposed into low and high-frequency components: the low-pass signals are blended linearly to smooth out illumination gradients, while high-pass signals utilize an abrupt transition to prevent “ghosting” or blurring of fine structural features. Tiles may optionally undergo CLAHE ^18^ pre-processing to homogenize brightness across the section.

The resulting full-resolution mosaics are then downsampled into thumbnails, one image file for each section. These low-resolution representations act as a computational bridge, serving as the primary input for the subsequent coarse thumbnail matching step. FEABAS includes an optional pre-downsampling high-pass filter. This step enhances biological membranes and edges at an intermediate resolution, ensuring that critical alignment features are preserved and legible even in the low-resolution thumbnail representation.

### Thumbnail Alignment

Following 2D stitching, FEABAS reconstructs the 3D volume through a multi-stage alignment process, beginning with the thumbnail matching step. The primary objective of thumbnail matching is to determine coarse correspondences between section pairs and provides the necessary initial constraints to guide subsequent high-resolution refinement. To prevent the gradual accumulation of errors (Z-drift) across large stacks, FEABAS registers sections beyond their immediate neighbors (e.g., matching section N against N±1, N±2, …).

The thumbnail matching workflow follows a two-step routine: (1) feature-based initialization via Radon transform features and RANSAC ^19^, followed by (2) dense refinement through grid-based template matching. The feature-based step is critical to the workflow’s success, as any unresolved rotations or large displacements are beyond the recovery range of localized template matching. Therefore, for challenging datasets involving fragmented or highly distorted sections, this initialization step employs a specialized iterative RANSAC strategy, rather than a single pass. In each iteration, RANSAC identifies the largest consistent match cluster among the remaining feature candidates. Then the system treats all matches from previous iterations as fixed constraints and performs an FEA relaxation on the new cluster. If the resulting residual errors are high, the system identifies the cluster as noise, rejects it as such and terminates the search. If the residuals are low, the matches are deemed consistent and merged into the global set. The area within the concave hull of these validated matches is then excluded from the candidate pool for subsequent iterations. This iterative approach forces the algorithm to systematically explore fragmented areas while ensuring every new match is physically compatible with the established global model. The success of the iterative RANSAC routine is also linked to the flexibility of the underlying finite element framework. In cases where tissue is physically discontinuous or folded, conventional elastic models often reject valid correspondences because the required non-smooth displacement generates prohibitive residual energy. To overcome this, FEABAS allows for the integration of coarse semantic masks that define regions of physical breakage. These masks indicate to the solver where the elastic continuum should be relaxed or severed, enabling the non-smooth deformations required to accommodate matches across fragmented sections.

The correspondences obtained from thumbnail matching are then integrated into a 3D FEA system (with very coarse meshes) encompassing the entire image stack to perform a rough global alignment. In this step, the system minimizes the total elastic energy across all sections simultaneously, effectively “pre-stacking” the volume. This global transformation provides a superior starting configuration for matching and optimization steps in the subsequent fine alignment stage.

### 3D Fine Alignment

The fine alignment stage produces the final registered volume by refining the rough correspondences established during the thumbnail phase. This process employs a significantly denser finite element mesh, typically with a nodal spacing of less than 5 μm, to capture localized elastic distortions across the stack. While the mesh domain for each section is automatically derived from the stitched montage boundaries by default, FEABAS also allows user-defined ‘material masks’ for complex cases. These masks supersede the automated geometry, enabling a semi-manual or manual yet precise discretization of physical discontinuities such as tears or folds. By locally modulating the mesh topology or stiffness, this mechanism enables the model to accurately represent tissue breakage or discontinuities, thereby maintaining smooth alignment in regions with compromised structural integrity.

Fine alignment matching operates at an intermediate isotropic resolution to find correspondences between neighbor sections. Empirically, using a resolution close to the physical section thickness provides an optimal balance between the accuracy and the computational cost. To initialize the dense meshes for the matched sections, the sparse matches and the transformations from the thumbnail stage are used to drive an FEA relaxation. Once initialized, the registration follows the Hierarchical Iterative Matching and Relaxation cycles described in the Algorithmic Foundation. This process alternates between grid-based template matching and mesh relaxation, progressively increasing the grid density to resolve fine-scale deformations.

With fine meshes reflecting the physical constraints of the section deformations generated and the dense pairwise correspondences established, FEABAS assembles these elements into a global 3D FEA system. Because the total degrees of freedom (DOFs) in a large-scale volume reconstruction can easily exceed the memory capacity of standard workstations, the software implemented two distinct strategies for the users to choose from to manage the computational load. The first strategy processes the volume in overlapping segments through a sequential sliding window. The solver selects a localized subset of sections and performs mesh relaxation to optimize their relative positions. Once a segment is resolved, the majority of its sections are fixed as a rigid reference for the subsequent window, while the sections at the trailing boundary remain flexible to mitigate fringe effects. The window then slides to the neighboring segment, propagating the alignment throughout the volume. This method is particularly valuable for workflows where alignment must occur concurrently with image acquisition, as it allows for the incremental integration of new sections into the existing stack without re-calculating the entire volume, but less parallelizable computationally. The alternative strategy utilizes a Hierarchical Chunk Optimization approach. The image stack is partitioned into independent blocks along the Z-axis, and a multi-step mesh relaxation procedure is performed. First, an intra-chunk relaxation resolves local distortions within each block independently. The solver then computes a convex hull for each aligned block to create a meta-section, treating the entire chunk as a single rigid entity. By utilizing the correspondences between sections at the interfaces between these meta-sections, the system aligns the meta-sections to form the global skeleton of the stack. These meta-section transformations are then applied back to the individual sections within each chunk, so that sections across chunks also align. Finally, the interior sections of each chunk are fixed, while targeted relaxation is applied at the interfaces to smooth transitions between chunks, ensuring a seamless volume without residual discontinuities from this block-wise process. This strategy is inherently more parallelizable, allowing independent chunks to be processed simultaneously across distributed workers or compute nodes, making it well-suited for high-throughput batch processing of large-scale datasets.

The flexibility of the FEM framework also enables efficient localized error correction. By leveraging Dirichlet boundary conditions, which allow specific nodal coordinates to be fixed as hard constraints, users can float only a problematic segment of the sections for re-optimization while keeping the surrounding volume anchored. This provides a surgical way to fix local alignment errors without affecting the global alignment.

In the final stage of the pipeline, the calculated transformations are applied to remap the full-resolution stitched sections into the reconstructed volume, as described in Image Rendering. The final volume is exported as either non-overlapping PNG tiles to be viewed in VAST ^20^, or a precomputed dataset for Neuroglancer ^21^.

## Result

### Real-World Application and Validation

The efficacy of FEABAS is established through its long-term deployment in production environments. Since its inception, the software (including its core algorithmic progenitors) has been utilized to process accumulatively more than one petabyte of electron microscopy data. To date, FEABAS has facilitated the published results of multiple works ^22–32^, and is the primary stitching and alignment tool for several forthcoming large-scale connectome studies. These high-throughput field tests have refined the tool into a robust, production-ready pipeline capable of handling the unpredictability of vEM datasets at scale.

To provide a benchmark against established state-of-the-art results, we utilized FEABAS to attempt reconstruction of the H01^33^ (human cortex) and MICrONS^34^ (mouse visual cortex) datasets. The original reconstructions of these landmark volumes represented a heroic effort, utilizing highly sophisticated and custom-built pipelines to overcome the unprecedented technical and computational challenges. Due to the immense computational and storage costs associated with archive retrieval and processing of petabytes-scale volumes in their entirety, we evaluated FEABAS on subsets of these datasets. For MICrONS, the test subset (provided by N. de Costa) covered 8.0 TB of raw data shared among 100 consecutive sections; and for H01, we processed 144.7 TB raw data making up 302 consecutive sections. Our smaller scale tests show that the physics-informed constraints of FEABAS can achieve at least comparable results with excellent axial continuity. To quantify registration accuracy, we employed Chunked Pearson’s Correlation (CPC), a benchmarking metric adopted in the MICrONS alignment study^7^. Aligned volumes were partitioned into 64 × 64 pixel chunks at a 32 nm resolution, and the Pearson correlation coefficient was calculated between corresponding chunks of adjacent sections. Regions devoid of tissue or obscured by imaging artifacts were excluded from the analysis. Benchmarking against published vEM volumes revealed that our pipeline achieved marginally higher CPC scores than the MICrONS dataset, with a more pronounced improvement observed over the H01 dataset (Figure 2a).

**Figure 2:**
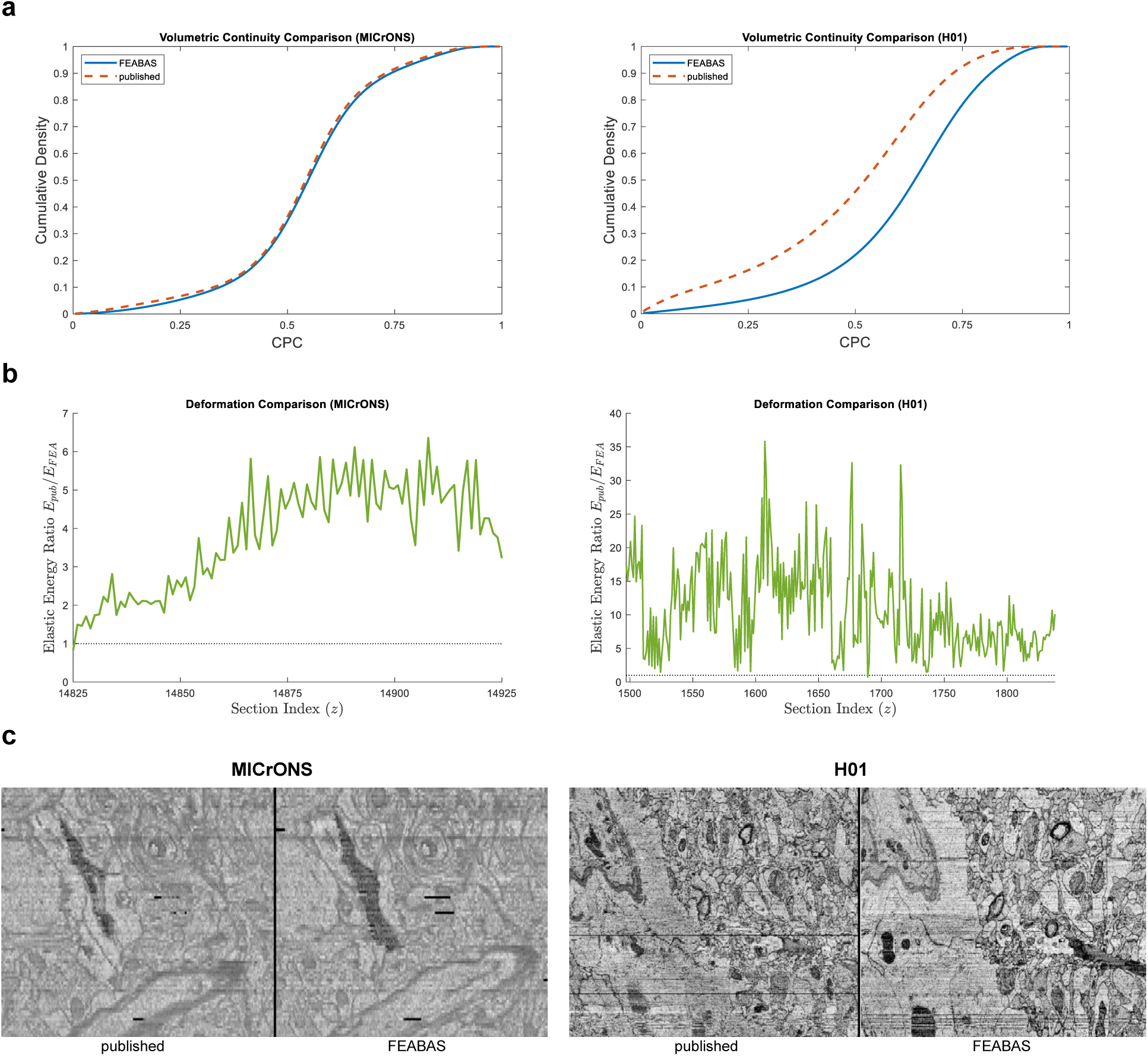
Quantitative and Qualitative Validation of Alignment Quality. **(a)** Volumetric Continuity Comparisons. Cumulative Distribution Function (CDF) of the Chunked Pearson Correlation (CPC) for the MICrONS (left) and H01 (right) datasets. In both cases, FEABAS (blue) exhibits a rightward shift compared to the published alignment ^33,34^(orange), indicating a higher proportion of high-correlation chunks. **(b)** Relative Geometric Deformation. Plot of the elastic energy ratio *E*_publised_/*E*_FEABAS_ across the section index (*z*) for MICrONS (left) and H01 (right). Values consistently greater than 1.0 demonstrate that FEABAS requires significantly less internal strain to satisfy registration constraints than the published method. This lower energy profile indicates that FEABAS achieves volumetric continuity with minimal non-physical warping, preserving the intrinsic biological geometry of the tissue more effectively. **(c)** Comparative Cross-Sectional Morphology. Representative *xz*-plane virtual cross-sections from the MICrONS (left) and H01 (right) volumes. Note: Gaps corresponding to missing sections were removed from the cross-sections for visual clarity. For a comprehensive, high-resolution comparison of both datasets, interactive Neuroglancer views are provided at https://lichtman.rc.fas.harvard.edu/feabas.

In addition to registration accuracy, the degree of induced deformation is a critical metric for evaluating alignment quality; excessive warping can improve axial continuity at the cost of distorting biological morphology or “verticalizing” the volume. An ideal alignment achieves high registration precision with minimal geometric strain. To compare the deformation required by our pipeline against published volumes (H01 and MICrONS), we estimated the underlying transformations by registering our FEABAS-montaged sections to their corresponding slices in the published datasets at 32 nm resolution. For consistency, we applied this same estimation procedure to our own aligned volume. We visually verified the registration accuracy for both our data and the published data to ensure the estimated transformations faithfully represented the actual alignments. We quantified the deformation for each section using the elastic energy with quadratic form ***u***^**T**^**K*u***, where **K** denotes the stiffness matrix and ***u*** represents the non-linear components of the mesh displacement. Our method achieved registration with a fraction of the geometric distortion seen in published volumes, which required on average 5.8 times (H01) and 3.6 times (MICrONS) more elastic energy than our approach (Figure 2b). While the deformation for the published MICrONS volume initially matched our results at one end of the z-range, its energy rapidly diverged. This initial similarity is likely artifactual, stemming from the use of as static “seed” references, given the fact that the folds in the first section of the published volume were not corrected. Because a proper restoration of biological continuity requires stretching these folds, leaving them uncorrected artificially deflates early deformation metrics before the distortion compounds in subsequent layers.

Interactive Neuroglancer links (https://lichtman.rc.fas.harvard.edu/feabas) are provided to allow direct visual comparison of our aligned volumes with the published datasets.

Beyond its use in the original developer’s environment, FEABAS has matured through open-source community engagement. Active use by independent research groups is evidenced by ongoing GitHub activity, where external users have contributed to the software’s robustness through bug reports and feature inquiries. This external validation ensures that the pipeline is not a “specialized” tool requiring the author’s intervention, but a general-purpose utility capable of being deployed on diverse, third-party datasets.

### Data Processing Rate

The processing throughput of FEABAS is influenced by dataset-specific characteristics, such as the density of artifacts, as well as the chosen mesh grid size, and hardware architecture. To provide a representative estimate of performance in practical applications, we benchmarked the pipeline using subsets of the H01 and MICrONS datasets with production-grade configurations. To demonstrate the platform agnosticism of the framework, the H01 data was processed on Google Cloud C2 virtual machines, while the MICrONS data was processed on the Harvard FASRC HPC cluster utilizing Intel Sapphire Rapids CPUs. During these runs, we utilized a dynamic range of 100 to 500 CPU physical cores, depending on the specific task and real-time resource availability. Because the majority of FEABAS operations are decoupled at the tile or section level, the system exhibits near-linear scaling with additional compute resources. To provide a universal metric for researchers planning similar scales of acquisition, we report our findings in (Table 1) as the number of CPU cores required to sustain a raw data throughput of 100 MB/s. Lower throughput in the MICrONS pipeline is a result of higher artifact density, requiring finer meshes compared to H01 to maintain alignment accuracy.

**Table 1:** Number of CPU Core Requirements per 100 MB/s Throughput.

|  |  | H01 | MICrONS |
| --- | --- | --- | --- |
| 2D stitching | matching | 29.72 | 21.31 |
|  | optimization | 0.25 | 0.39 |
|  | render | 54.36 | 62.17 |
| thumbnail alignment | downsample | 0.13 | 0.92 |
|  | matching | 0.14 | 0.06 |
| 3D fine alignment | matching | 17.72 | 47.40 |
|  | optimization | 0.16 | 1.08 |
|  | render | 30.90 | 52.85 |
| total pipeline |  | 133.38 | 186.18 |

Note that the reported throughput figures (Table 1) cover the core registration and rendering pipeline; upstream artifact mask generation is excluded, as this is treated as a decoupled preprocessing step that can be handled by various external semantic segmentation tools. Based on these standardized requirements, we estimate that the continuous image acquisition rates originally reported for the H01 (190MB/sec ^33^) and MICrONS (2PB in 6 months ^34^) studies would require approximately 250 and 240 CPU cores, respectively, to maintain real-time processing.

### Elastic Stitching Yields Seamless Montages

Intrinsic distortions in vEM acquisition, ranging from systematic lens aberrations to stochastic charging effects, impose non-linear deformations that a standard affine transformation cannot fully resolve. In these cases, restricting the alignment to affine parameters is insufficient, often leaving visible seams and misalignments at tile boundaries. We demonstrate that an elastic stitching approach is required to achieve sub-pixel continuity, as evidenced by the MICrONS dataset where affine-only montages exhibit prominent stitching artifacts that are seamlessly resolved through elastic relaxation (Figure 3a-c).

**Figure 3:**
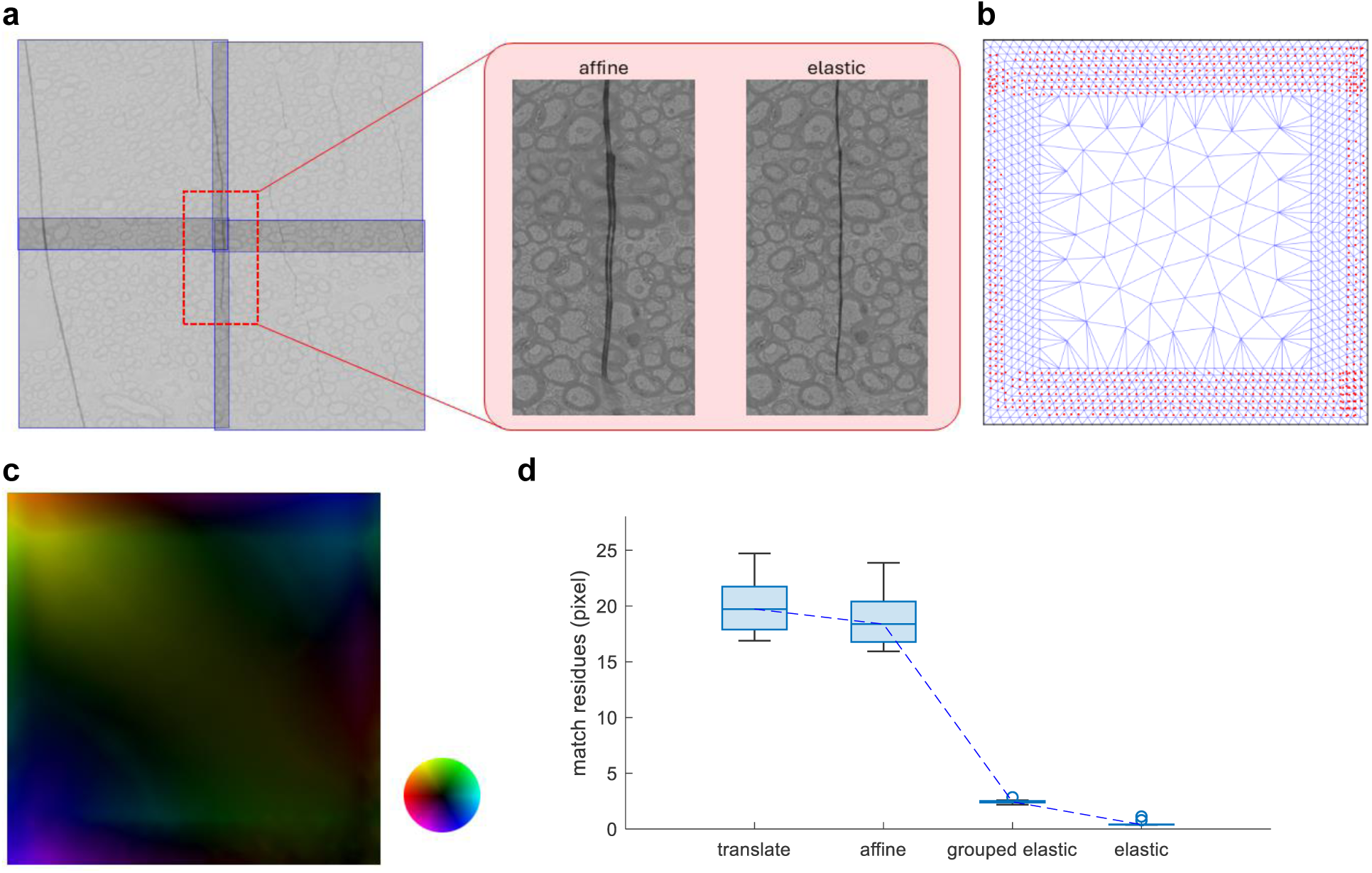
Elastic 2D Stitching and Adaptive Mesh Relaxation. **(a)** Comparison of Stitching models. Left: A 2×2 montage of semi-transparent image tiles, cropped from MICrONS dataset, with marked boundaries (blue), illustrating the overlapping regions. Center/Right: Magnified views of the region indicated by the red rectangle. The affine-only reconstruction (center) reveals seam artifacts at tile interfaces. In contrast, the elastic stitching result (right) demonstrates the seamless restoration of feature continuity across boundaries. **(b)** Adaptive Mesh Configuration and Landmark Distribution. Visualization of the finite-element mesh used for the elastic stitching shown in a. The mesh density is selectively increased at tile interfaces (borders) to accommodate the higher degrees of freedom required for resolving seam errors. Red dots indicate the distribution of landmark correspondences established via coarse-to-fine template matching; these matched correspondences serve as the driving force in the mesh relaxation process. **(c)** Elastic Deformation Field. Mapping of the non-linear displacement field required for seamless stitching in a MICrONS tile. Hue represents displacement direction and brightness indicates magnitude (normalized such that the maximum brightness corresponds to a 25-pixel displacement), as defined by the inset color wheel. The observed pattern illustrates the non-affine nature of the underlying distortions. **(d)** Distribution of residual Euclidean distances between matched landmarks across successive optimization stages: translation, affine, grouped elastic, and localized elastic relaxation. Boxplots show the reduction in error at each stage, with the largest decrease occurring during the grouped elastic phase.

Quantitative analysis of the residual displacement between matches illustrates the progression of this refinement. In the MICrONS dataset, we observed that the most substantial reduction in residual error occurs after grouped elastic relaxation (Figure 3d), which specifically compensates for the systematic lens distortion of the electron-optical system. While this suggests that lens aberration is the primary driver of misalignment in this specific dataset, the final localized elastic step remains essential to bring down the remaining residuals caused by non-systematic distortions. Although not suitable for our benchmark datasets, FEABAS provides users with the flexibility to navigate this hierarchy; if the residual drop after the affine stage meets the project’s requirements, the elastic steps can be bypassed to prioritize throughput.

### Correcting Mechanical Artifacts During Cross-Section Registration

For large-scale structural damage, such as the broken sections present in the mouse cortex dataset ^32^, the thumbnail alignment pipeline can automatically reassemble disjointed tissue. By applying material masks to demarcate individual tissue fragments and exclude interstitial gaps, the separated pieces are permitted to translate and deform independently. As shown in Figure 4a-e, FEABAS successfully matches these isolated fragments to their corresponding locations in the adjacent section, effectively reconstructing the broken tissue within the unified global coordinate frame.

**Figure 4:**
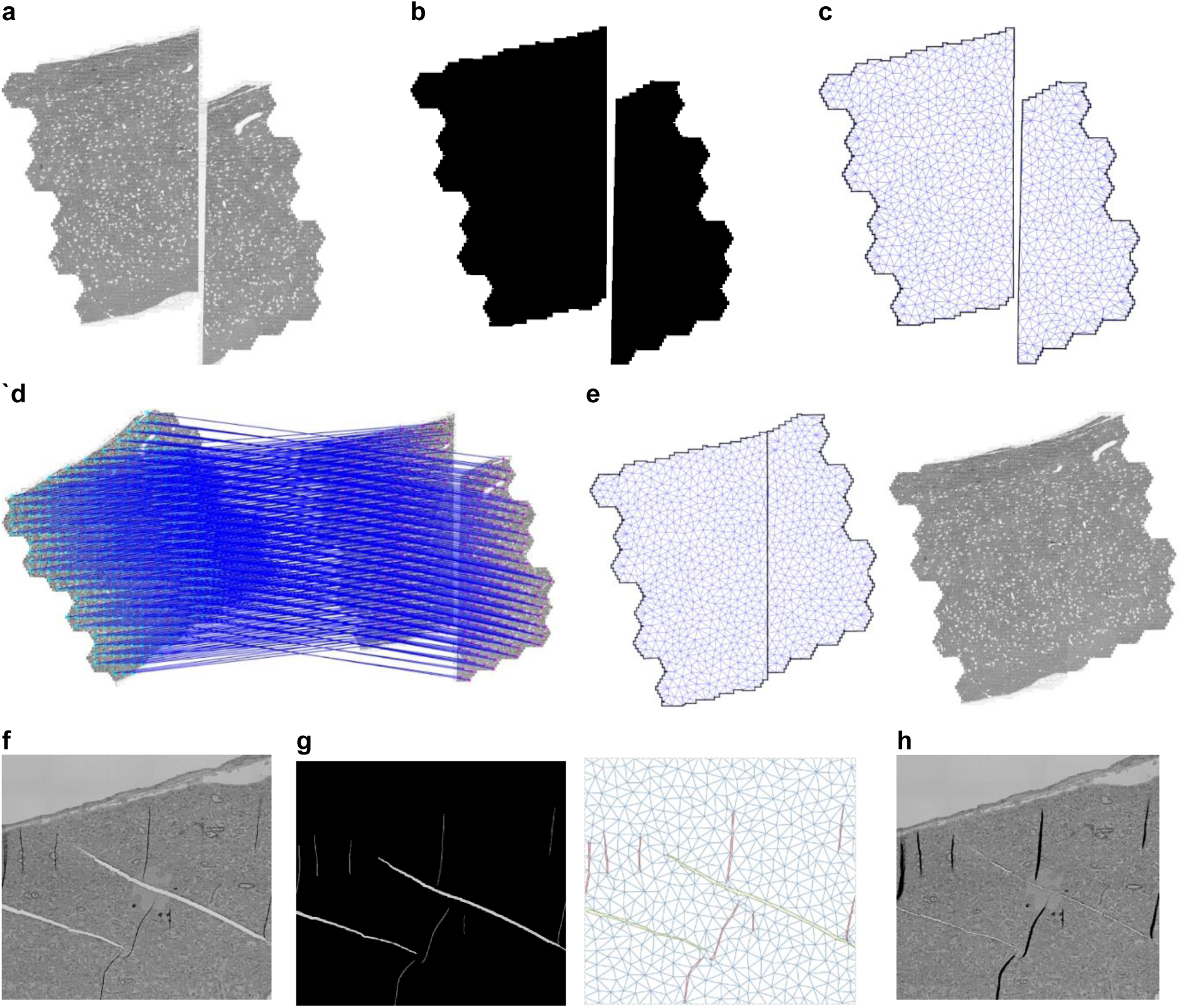
Artifacts Correction During 3D alignment. **(a–e)** Global Thumbnail Alignment of Fragmented Tissue. **(a)** A mouse cortex^32^ section thumbnail featuring a major discontinuity where the tissue has split into two disjointed fragments. **(b)** The corresponding material mask input to the workflow; black regions indicate valid tissue. **(c)** The generated finite-element mesh, where stiffness is assigned exclusively to the intact tissue regions defined in b. **(d)** Landmark correspondences (blue lines) established between the fragmented section and a neighboring reference. **(e)** The reconstructed section post-alignment, shown as both a transformed mesh and rendered image, demonstrating the restoration of global tissue macro-geometry. **(f–h)** Localized Resolution of Folds and Cracks. **(f)** A high-resolution crop from the MICrONS dataset featuring compressed folds (dark) and micro-cracks (bright). **(g)** Micro-cracks (light gray in mask) are mapped to zero-stiffness elements (olive) to facilitate gap closure. Folds (dark gray in mask) are assigned anisotropic stiffness (red) that favors expansion over compression to revert the folding process. **(h)** The final aligned image, demonstrating the successful stretching of folds and closure of cracks to restore local biological continuity.

To address localized physical artifacts, FEABAS leverages its adaptive mesh to selectively apply distinct mechanical properties to identified defect boundaries. We tested the framework on the MICrONS dataset, which was heavily affected by folds and small cracks. As shown in Figure 4f-h, FEABAS effectively corrects these features: the solver expands compressed folds to their original surface area and collapses artificial gaps created by cracks to restore biological continuity.

## Discussion

### Rationale for the Finite Element Method

The fundamental advantage of the finite-element approach lies in its ability to model vEM data as a piecewise-smooth physical system rather than as a purely geometric image stack. While tissue sections generally exhibit smooth deformations, they are frequently punctuated by “kink points” (e.g. tears, folds) where the universal smoothness assumptions of interpolation-based ^1–3^ or regular-grid ^4–6^ methods fail. By utilizing an adaptive mesh, FEABAS can concentrate degrees of freedom around these critical features, capturing complex non-linear distortions while maintaining a sparse computational representation. While the utility of adaptive meshes for resolving EM artifacts has been explored previously^35^, FEABAS goes beyond geometric triangle fitting by adopting a principled physical model using FEA. This formulation provides a level of conceptual clarity, operational flexibility, and numerical robustness that is difficult to achieve with purely geometric treatments. Furthermore, the sparse FEA construction enables direct computation of high-fidelity transforms at full resolution with moderate computational cost. This bypasses the need to compute the saturated fields typical of deep learning approaches^7–9^.

Unlike other spring-mesh frameworks ^4,5,35^ that rely on transient simulations, which can be computationally slow and prone to local minima, FEABAS directly solves for the static equilibrium of the entire system. Mathematically, the FEA formulation reduces the equilibrium-finding problem to a sparse linear system, enabling the use of highly optimized, established solvers well beyond the scope of vEM. This global relaxation also allows treating each section equally and thereby naturally distributes residual errors across the Z-axis, preventing the over-registration and cumulative drift common in pipelines that register sections sequentially to a fixed reference frame.

Beyond its mathematical simplicity, the FEA paradigm offers a level of interpretability that black-box neural networks cannot match. For the researcher, the model’s physical nature provides clear intuition: alignment failures can be diagnosed and corrected through the lens of stress, strain, and material stiffness. It replaces the unpredictable behavior of automated black-box systems with a robust and transparent foundation for large-scale connectomics.

A notable operational requirement of the FEA approach is the need for material masks to identify artifact locations. Unlike end-to-end deep learning (DL) methods that implicitly learn these features, FEABAS requires explicit spatial context to define the mesh’s behavior. It is worth noting, however, that even some DL-based registration frameworks have moved toward using defect masks as auxiliary inputs ^7^ to improve reliability, suggesting that explicit artifact handling is often necessary regardless of the underlying solver. In practice, the effort required to generate these masks depends on the dataset’s scale and quality. For volumes with localized defects, manual annotation remains a viable and high-precision option. For larger-scale datasets where manual intervention becomes a bottleneck, the masking process can be delegated to automated DL-based segmentation tools^11^. We acknowledge that generating masks can be time-consuming, especially for artifacts with subtle appearances, where automated detection can fail. Nevertheless, we argue that decoupling semantic interpretation (masking) from geometric registration (alignment) offers a strategic advantage. By isolating artifact detection as a standard semantic segmentation task, FEABAS allows researchers to employ the most current and accessible DL models independently of the specialized alignment solver. This modularity ensures that the registration remains transparent and interpretable, as researchers can directly inspect and refine the masks that govern the solver’s physics.

### Mitigation of Z-Axis Alignment Drift

A pervasive challenge in vEM reconstruction is alignment drift, a phenomenon where consecutive sections appear locally continuous, but the volume exhibits a slow, low-frequency lateral wobble when navigating along the z-axis. Correcting drift is notoriously difficult due to the general absence of volumetric ground truth. In our experience, this drift primarily originates from two distinct sources: stochastic matching error accumulation and systematic physical deformation.

The first source is the accumulation of slight inaccuracies during matching steps. As alignment propagates through a large stack, minute pairwise errors can compound in a random-walk fashion, gradually pulling the trajectory off-center. This form of drift is relatively straightforward to mitigate computationally; users can increase the resolution of the feature matching step or introduce long-distance (skip-section) matching pairs to provide global geometric constraints that anchor the random walk.

The second, more intractable source of drift arises from inherent, systematic changes in section deformation. For example, replacing a diamond knife mid-run can abruptly alter the tissue’s compression rate. The FEABAS spring-mesh system is inherently designed to distribute deformations evenly along the z-axis, a practice that works exceptionally well under the assumption of consistent or random local distortions. The current default alignment settings, particularly the mesh stiffness and relaxation process, are empirically tuned to prioritize Z-axis consistency while minimizing excessive deformation under such an assumption, and were used on the two benchmark datasets in this work. However, when faced with an abrupt, systematic shift in the physical data, the mesh’s uniform elastic resistance resists the sudden change, inadvertently distributing the corrective deformation across many adjacent sections and manifesting as pronounced drift. To remedy this physically induced drift, FEABAS allows users to modulate the mesh mechanics. First, the global stiffness of the mesh can be reduced (e.g., to 0.05x the default), thereby loosening the regularization penalty and permitting the large-scale deformations required to bridge the knife change. Second, we employ an iterative “annealing” technique: the aligned transformation from the first pass is adopted as the new zero-strain resting state for subsequent rounds of relaxation. This iteratively diminishes the mesh’s elastic resistance to true physical changes. As demonstrated in Supplementary Figure 2, while default stiffness settings can result in severe lateral drift around a knife-change boundary, applying a softer mesh noticeably smooths the transition, and subsequent rounds of annealing almost eliminate the drift.

Looking forward, a more definitive solution to alignment drift likely lies outside of purely serial 2D image registration. When available, undistorted 3D reference data, such as micro-CT scans of the embedded tissue block prior to sectioning, could provide an absolute structural ground truth. We speculate that using a coarse registration to a micro-CT volume to establish the initial resting state for the high-resolution FEABAS elastic solver could theoretically eliminate z-drift from the outset, bridging the gap between global biological morphology and ultrastructural continuity.

## Code Availability

The FEABAS software, default configuration files, and documentation are open-source and publicly available on GitHub at https://github.com/YuelongWu/feabas.

## Acknowledgements

The authors thank the Connectomics team at Google, specifically Tim Blakey, Michał Januszewski, and Viren Jain, for providing access to the H01 raw dataset and the Google Cloud Platform (GCP) computational resources that facilitated large-scale processing. We are grateful to Nuno Maçarico da Costa and Russell Torres (Allen Institute for Brain Science), and H. Sebastian Seung (Princeton University) for providing the raw data and support for the MICrONS dataset. We thank Mariela Petkova and the Engert Lab (Harvard University) for permission to include “Fish 2.0” datasets ahead of their primary publication. We also acknowledge the FEABAS open-source community for their invaluable feedback, bug reports, and contributions to the software’s development. Finally, the lead author thanks little Cora for all the inspiration and joy during the final stage of this work.

This work was supported by the National Institutes of Health (NIH) grants U19NS104653, UG3MH123386, U24NS109102, RM1NS132981, UM1NS132358, and R01NS133654; the National Science Foundation (NSF) grant IOS-2227964; and the Office of Naval Research (ONR) grant N00014-20-1-2828.

---

**Supplementary Figure 1:**
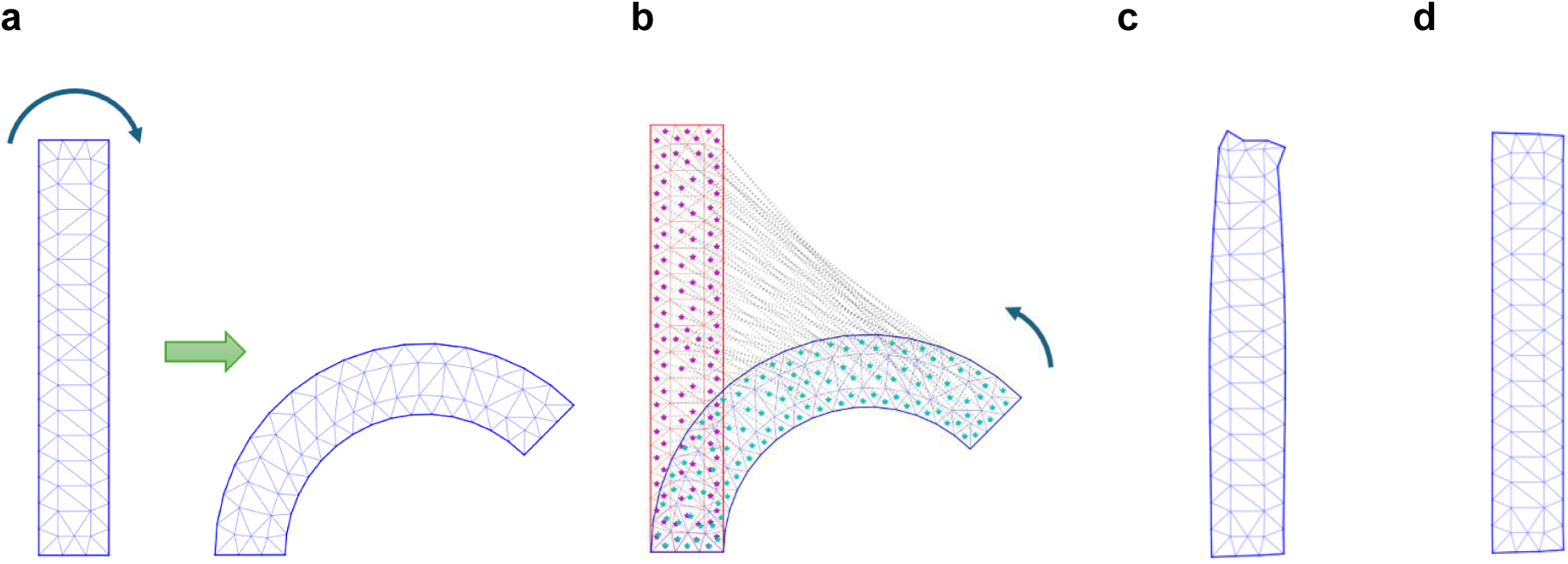
Demonstration of Rotation-Variant (Linear) vs. Rotation-Invariant (hyperelastic) Elasticity Models. **(a)** Geometric setup of the elastic stress test. An undeformed rectangular reference mesh (left) is manually bent into an arch-like geometry (right), introducing a severe, non-uniform rotation gradient from one end to the other end of the structure. **(b)** Landmark constraint configuration. To reverse the deformation, landmark correspondences (asterisks) are established at the centroid of every triangular element, mapping points from the deformed target mesh (blue) back to their original coordinates in the undeformed reference configuration (red). **(c)** Optimization result using a standard linear elasticity model. Elements at the highly rotated end (top) exhibit severe non-physical distortions, due to the rotation-variant nature of the model. **(d)** Optimization result using the nonlinear St. Venant-Kirchhoff (SVK) hyperelastic model. By accounting for finite strains, the SVK model accommodates the large rotation without severe element distortions.

**Supplementary Figure 2:**
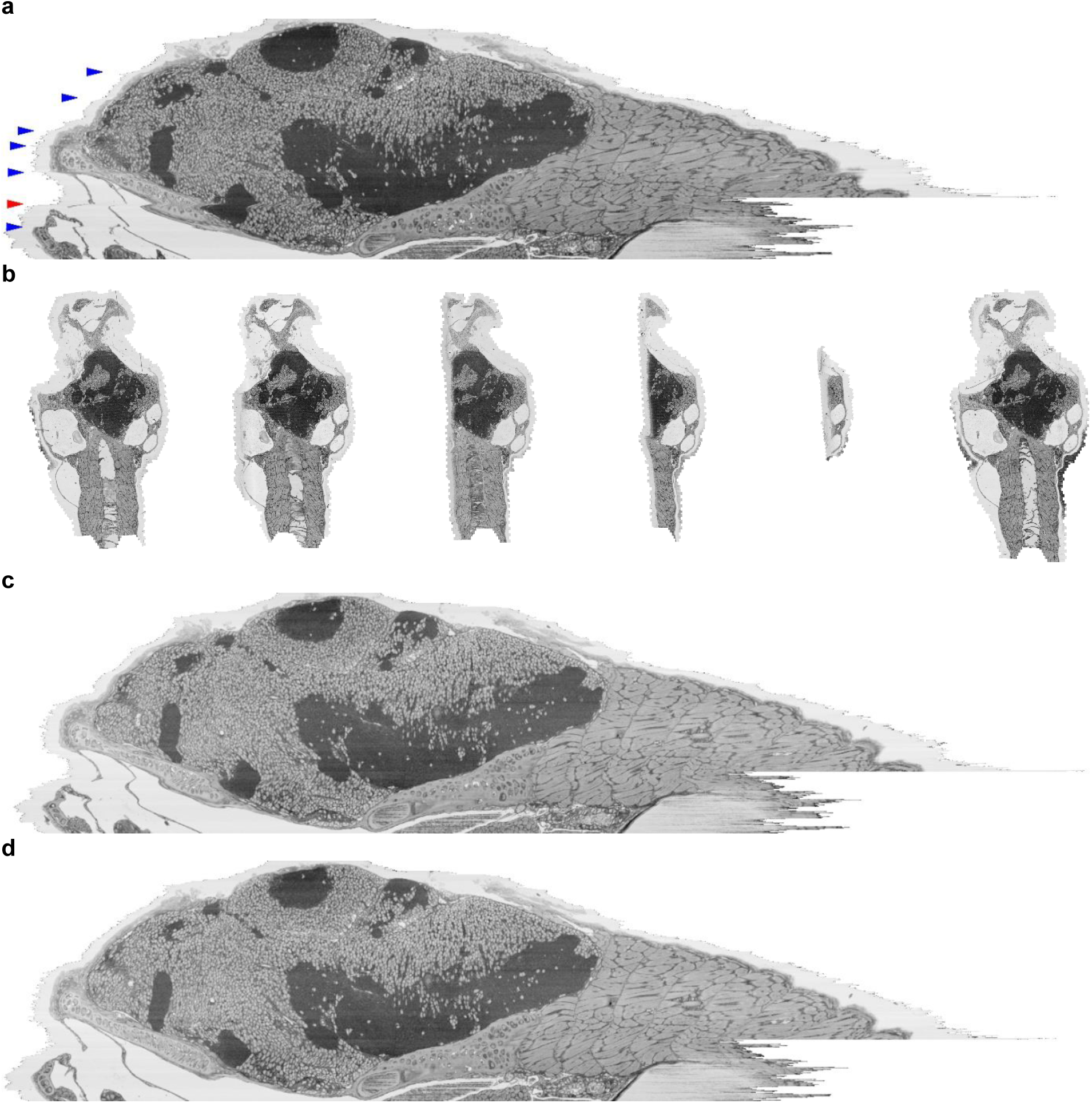
Mitigation of Volumetric Z-Drift. **(a)** Virtual xz cross-section of the zebrafish dataset processed with default mesh stiffness. Significant axial drift is visible (arrows). The red arrow denotes the onset of the most severe discontinuity. **(b)** Serial thumbnails of the sections corresponding to the red arrow in a. The sequence illustrates a gradual transition from complete sections to partial fragments and back to complete sections, a hallmark of diamond knife replacement. **(c)** The same xz view as in a, processed with the mesh stiffness reduced to 0.05x the default value. The reduction in mechanical resistance allows the solver to better accommodate the geometric step-change, partially attenuating the observed drift. **(d)** xz reconstruction following three rounds of “annealing.” In this process, the relaxed mesh state from each iteration is defined as the new resting position for the subsequent solve. This iterative realignment effectively eliminates the remaining drift.

## References

1. Cardona, A. et al. TrakEM2 software for neural circuit reconstruction. PLoS One 7, e38011 (2012).

2. Khairy, K., Denisov, G. & Saalfeld, S. Joint Deformable Registration of Large EM Image Volumes: A Matrix Solver Approach. arXiv [cs.CV] (2018).

3. Watkins, P. V., Jelli, E. & Briggman, K. L. msemalign: a pipeline for serial section multibeam scanning electron microscopy volume alignment. Front. Neurosci. 17, 1281098 (2023).

4. Saalfeld, S., Fetter, R., Cardona, A. & Tomancak, P. Elastic volume reconstruction from series of ultra-thin microscopy sections. Nat. Methods 9, 717–720 (2012).

5. Suissa-Peleg, A. Mb_aligner. (Github, 2019).

6. Wetzel, A. W. et al. Registering large volume serial-section electron microscopy image sets for neural circuit reconstruction using FFT signal whitening. in 2016 IEEE Applied Imagery Pattern Recognition Workshop (AIPR) 1–10 (IEEE, 2016).

7. Popovych, S. et al. Petascale pipeline for precise alignment of images from serial section electron microscopy. Nat. Commun. 15, 1–15 (2024).

8. Yoo, I., Hildebrand, D. G. C., Tobin, W. F., Lee, W.-C. A. & Jeong, W.-K. ssEMnet: Serial-Section Electron Microscopy Image Registration Using a Spatial Transformer Network with Learned Features. in Deep Learning in Medical Image Analysis and Multimodal Learning for Clinical Decision Support 249–257 (Springer International Publishing, 2017).

9. Xin, T. et al. A novel registration method for long-serial section images of EM with a serial split technique based on unsupervised optical flow network. Bioinformatics 39, (2023).

10. Shewchuk, J. R. Delaunay refinement algorithms for triangular mesh generation. Comput. Geom. 22, 21–74 (2002).

11. Lin, Z., Wei, D., Lichtman, J. & Pfister, H. PyTorch connectomics: a scalable and flexible segmentation framework for EM connectomics. arXiv preprint arXiv:*2112.05754* (2021).

12. Bell, N., Olson, L. N. & Schroder, J. PyAMG: Algebraic Multigrid Solvers in Python. J. Open Source Softw. 7, 4142 (2022).

13. Bonet, J. & Wood, R. D. Nonlinear Continuum Mechanics for Finite Element Analysis. (Cambridge University Press, Cambridge, England, 2010). doi:10.1017/cbo9780511755446.

14. Berg, M., Cheong, O., Kreveld, M. & Overmars, M. Point Location. in Computational Geometry, Algorithms and Applications 121–146 (Springer Berlin Heidelberg, 2008).

15. Duchon, C. E. Lanczos Filtering in One and Two Dimensions. J. Appl. Meteorol. Climatol. 18, 1016–1022 (1979).

16. Hunter, J. D. Matplotlib: A 2D Graphics Environment. Comput. Sci. Eng. 9, 90–95 (2007).

17. Eberle, A. L. & Zeidler, D. Multi-Beam Scanning Electron Microscopy for High-Throughput Imaging in Connectomics Research. Front. Neuroanat. 12, 112 (2018).

18. Pizer, S. M. et al. Adaptive histogram equalization and its variations. Computer Vision, Graphics, and Image Processing 39, 355–368 (1987).

19. Fischler, M. A. & Bolles, R. C. Random sample consensus: a paradigm for model fitting with applications to image analysis and automated cartography. Commun. ACM 24, 381–395 (1981).

20. Berger, D. R., Seung, H. S. & Lichtman, J. W. VAST (Volume Annotation and Segmentation Tool): Efficient Manual and Semi-Automatic Labeling of Large 3D Image Stacks. Front. Neural Circuits 12, 88 (2018).

21. Maitin-Shepard, J. Neuroglancer: WebGL-Based Viewer for Volumetric Data. (Github, 2024).

22. Karlupia, N. et al. Immersion fixation and staining of multicubic millimeter volumes for electron microscopy-based connectomics of human brain biopsies. Biol. Psychiatry 94, 352–360 (2023).

23. Lu, X., et al. A scalable staining strategy for whole-brain connectomics. bioRxivorg (2023) doi:10.1101/2023.09.26.558265.

24. Bidel, F. et al. Connectomics of the Octopus vulgaris vertical lobe provides insight into conserved and novel principles of a memory acquisition network. Elife 12, (2023).

25. Sant, H., et al. 3D reconstructions of peptidergic terminals suggest synaptic pruning in an adult molluscan brain. 65, S457–S457 (2025).

26. Meirovitch, Y., et al. SmartEM: machine-learning guided electron microscopy. bioRxivorg 2023.10. 05.561103 (2024) doi:10.1101/2023.10.05.561103.

27. Boulanger-Weill, J., et al. Correlative light and electron microscopy reveals the fine circuit structure underlying evidence accumulation in larval zebrafish. bioRxivorg (2025) doi:10.1101/2025.03.14.643363.

28. Zhang, S. et al. Ultrastructural reconstruction of the endodermal nerve net of Hydra vulgaris. Curr. Biol. 35, 5490–5501.e3 (2025).

29. Perks, K. E., et al. Connectome analysis of a cerebellum-like circuit for sensory prediction. bioRxivorg (2025) doi:10.1101/2025.07.03.662989.

30. Petkova, M. D., et al. A connectomic resource for neural cataloguing and circuit dissection of the larval zebrafish brain. bioRxivorg (2025) doi:10.1101/2025.06.10.658982.

31. Tian, Q. et al. Quantifying axonal features of human superficial white matter from three-dimensional multibeam serial electron microscopy data assisted by deep learning. Neuroimage 313, 121212 (2025).

32. Lu, X., et al. Probing molecular diversity and ultrastructure of brain cells with fluorescent aptamers. bioRxivorg (2023) doi:10.1101/2023.09.18.558240.

33. Shapson-Coe, A. et al. A petavoxel fragment of human cerebral cortex reconstructed at nanoscale resolution. Science 384, eadk4858 (2024).

34. MICrONS Consortium. Functional connectomics spanning multiple areas of mouse visual cortex. Nature 640, 435–447 (2025).

35. Scheffer, L. K., Karsh, B. & Vitaladevun, S. Automated Alignment of Imperfect EM Images for Neural Reconstruction. arXiv [q-bio.QM] (2013).

36. Januszewski, M. Sofima: Scalable Optical Flow-Based Image Montaging and Alignment. (Github, 2022).

